# Environmental and temporal factors affecting record white-tailed deer antler characteristics in Ontario, Canada

**DOI:** 10.1101/2025.03.25.645257

**Authors:** Brooklyn S. Cars, Joseph M. Northrup, Keith Beasley, Paul Beasley, Kevin Beasley, Aaron B.A Shafer

**Affiliations:** Environmental and Life Sciences Graduate Program, Trent University, 2140 East Bank Drive, Peterborough, ON K9J 7B8, Canada; Environmental and Life Sciences Graduate Program, Trent University, 2140 East Bank Drive, Peterborough, ON K9J 7B8, Canada and Wildlife Research and Monitoring Section, Ontario Ministry of Natural Resources, Peterborough, Ontario, Canada; The Foundation for the Recognition of Ontario Wildlife, Ontario, Canada

**Keywords:** white-tailed deer, *Odocoileus virginianus*, antler size, tine configuration, environmental factors, Ontario, hunters, harvest, climate effects, land cover

## Abstract

White-tailed deer (*Odocoileus virginianus*) are an ecologically and economically important species in North America. Their antlers, one of their most recognizable features, are used for dominance displays, mate attraction, and defense, with size and shape being key determinants of success. Antler characteristics are influenced by a combination of genetics, age and environmental factors, notably habitat quality and resource availability. In this study, we explored how diverse environmental factors, including climate and land cover composition, impact antler size, tine configuration, and the distribution of record-scoring white-tailed deer across Ontario, Canada, using hunter-submitted data from long-term antler scoring records. We used conditional autoregressive (CAR) models to examine these relationships and found that warmer temperatures the year of harvest were positively associated with larger antlers and more record deer in a given county, while winter precipitation the year of harvest was negatively associated with these characteristics, likely due to reduced forage availability or increased energy expenditure during more severe winters. Rangeland and forest land cover types were positively associated with increased antler size and tine number. We observed no temporal changes in antler size in Ontario, contrasting with broader trends observed in North America. These results show how local environmental conditions and land cover composition influence antler traits and the distribution of record white-tailed deer, highlighting the complexity of environmental influences on trait variation.

---

The Cervidae (deer) family emerged during the late Miocene where they developed the secondary sex characteristics that we associate with modern deer (Gilbert et al. 2006). One such secondary sex characteristic, antlers, are bony extensions of the skull that generally shed every year, and apart from caribou (*Rangifer tarandus*) are typically only found on males (Rue 1989, Hewitt 2011). Antlers have multiple purposes including dominance displays and interactions, weapons against predators and other male Cervidae, and indicators used in mate selection (Clutton-Brock 1982, Goss 1983, Morina et al. 2018). In white-tailed deer (*Odocoileus virginianus*), males are the dominant sex where social dominance is largely dictated by age and body mass, both of which are closely tied to antler size (Townsend and Bailey 1981). Fully mature males tend to have larger antlers due to the lower energy demands for body growth and thus their ability to allocate more resources (Goss, 1983). Likewise, in red deer (*Cervus elaphus*), larger antlers are associated with increased reproductive success (Kruuk et al. 2002). But while larger-antlered males may have an advantage when it comes to breeding opportunities, mating success is not limited to them (DeYoung et al. 2006). Younger or smaller-antlered white-tailed deer can achieve mating success if they are more dominant and in good overall health, in populations with a female-skewed sex ratio or a younger male age structure, or by avoiding direct competition with larger males and engaging in opportunistic mating (Townsend and Bailey 1981, DeYoung et al. 2006, Turner et al. 2016).

Although antler traits have a heritable component (e.g. Michel *et al*., 2016; Jamieson *et al*., 2020), environmental conditions and habitat composition affect their expression, with the quality and quantity of forage among landcover types playing a large role. Agricultural and rangeland areas provide nutrient rich, easily accessible forage, including plants such as soybeans, alfalfa, and clover (Harper 2019), while forests offer cover from predators and winter conditions but the quality of forage varies with tree species and understory composition (Voigt et al. 1997). Consistent with these patterns, Strickland and Demarais (2008) observed a positive association in the southern United States between mean antler size and landcover types that favour forage-producing vegetation such as agriculture. Further, Cain et al. (2019) showed that larger-antlered deer in the Midwest United States were associated with an intermixed landscape with more fragmented patches of land cover, suggesting that habitat composition and variety can shape antler growth through changes in resource availability. In addition to landcover, seasonal environmental conditions such as heavy winter snowfall can reduce forage availability and increase energetic costs for white-tailed deer populations. For example, Garroway and Broders (2005) found that deer in areas with deeper snow exhibited lower body condition, suggesting that winter severity directly affects white-tailed deer energy reserves.

Intergenerational effects can also influence antler and other physical traits (Freeman et al. 2013), with, for example, maternal nutrition influencing offspring body mass and antler size in white-tailed deer (Monteith et al. 2009). Strickland et al. (2020) showed that summer droughts during gestation and early lactation caused maternal nutritional stress that limited resource allocation to developing offspring and resulted in reduced antler mass in young males. Similarly, Peterson et al. (2019) found that male white-tailed deer born during years with extreme droughts had smaller antler metrics compared with those born in non-extreme years. Collectively, these studies highlight the need to consider multi-year and spatial environmental covariates when modelling trait variation.

Across North America, hunter harvest is the primary means of managing white-tailed deer and management strategies vary widely, ranging from highly regulated systems with antler□point restrictions and controlled harvests (Strickland and Demarais 2007) to more liberal frameworks with longer seasons and fewer restrictions on age or sex class (Florida Fish And Wildlife Conservation Commission 2025). In Ontario, Canada, the hunting system leans toward a moderately liberal framework; hunters can purchase a white-tailed deer licence and an antlered deer (i.e. buck) tag valid for any open deer season, with the option to purchase additional tags depending on the management unit. Ontario regulations define an antlered deer as one with at least one antler measuring 7.5 cm or more in length, which limits harvest of younger or smaller-antlered bucks helping preserve future breeding males. Antlerless deer (female/fawn buck) tags are awarded via random draw for a specific Wildlife Management Unit (WMU) which allow wildlife managers to regulate population growth by controlling female harvest (Government of Ontario 2019*a*). Both residents and non-residents can hunt in Ontario, with harvest levels fluctuating among WMUs based on deer density, land access, and hunter participation. Approximately 50,000–75,000 deer are harvested annually across the province (Government of Ontario 2025).

With the allocation of antlered deer tags being more liberal than antlerless tags, hunters have greater opportunity to harvest male white-tailed deer. Regardless of hunting motivation (e.g. food, trophy), hunters generally prefer larger bucks, including trophy bucks with larger antlers (D’Angelo and Grund 2015, Boone and Crockett Club 2025). With larger, higher-scoring bucks often experiencing heavier harvests, there has been a broadscale negative trend in the mean antler size of trophy white-tailed deer across North America over the last century, largely attributed to reductions in male age structure resulting from harvest pressure (Monteith et al. 2013). But while selective harvest can influence age structure, the effects on population-level antler size and mating structure are complex. Males of all ages and antler sizes typically breed successfully, and management strategies often allow younger males to reach prime antler size before harvest (Webb et al. 2012, Turner et al. 2016). Thus, the potential evolutionary impacts of removing larger males on mating structure and antler size are likely limited (Webb et al. 2012).

This study aims to determine which environmental factors influence antler size and tine configuration in white-tailed deer across Ontario, Canada. Long-term antler scoring records provided us with a unique opportunity to examine white-tailed deer antler characteristics and temporal patterns over time and space in the province. We examined the influence of environmental factors on the spatial distribution, tine number, antler symmetry, and overall antler score of record white-tailed deer across Ontario. We hypothesized that antler size in white-tailed deer is positively associated with environmental factors that promote forage availability and minimize energy expenditure during harsh winter conditions, such as adequate cover and reduced snow depth. Specifically, we predicted larger antlers in counties with higher proportions of agricultural, rangeland, and forest cover, which provide accessible food and shelter. We also hypothesized that warmer temperatures during the growing season would be associated with increased antler size, while higher winter precipitation would have a negative effect due to increased energetic demands. Finally, we hypothesized that selective harvest of large-antlered individuals over time has led to a temporal decline in antler size across Ontario through reductions in age structure.

## STUDY AREA

White-tailed deer in Ontario, Canada, occupy a diverse landscape with varying anthropogenic influences. Historically, white-tailed deer populations have been concentrated in southern Ontario, where winters are milder and growing seasons are longer compared to more northern parts of the province (Kennedy-Slaney et al. 2018). This area is dominated by agricultural and deciduous forests, which provide high-quality forage such as forbs, legumes, and other crops during spring and summer (Ecological Stratification Working Group 1995, Government of Ontario 2021). However, white-tailed deer exist throughout parts of central and northern Ontario, including the boreal forest region, where they experience harsher winters. Their summer habitats consist of early successional forests with abundant food sources which shifts to coniferous and mixed-wood forests in winter that offer shelter from excessive snowfall and winter browse (Voigt et al. 1997, Hewitt 2011). Though browse is lower in nutritional quality, it is readily used by white-tailed deer during Ontario’s colder seasons, making up 74-91% of their diet when other vegetation is scarce (Mautz *et al*., 1976; Gray and Servello, 1995; Hewitt, 2011). Where they overlap, wolves (*Canis lupus*) are the primary predators of white-tailed deer in Ontario, while coyotes (*Canis latrans*) fill that role in their absence, though do not fully replace wolves ecologically (Benson et al. 2017). In addition to climate variation, Ontario’s expanding infrastructure and growing population has tripled in the last 50 years (Government of Canada 2016, 2022), consequently reducing the available habitat and forage available for deer. As a result, deer are increasingly concentrated in the habitats that remain suitable, including forest patches, crop fields, and early successional areas (Voigt et al. 1997, Chen et al. 2015). However, the proliferation of agricultural landscapes might increase food availability in some regions further north (Hewitt 2011).

## METHODS

### Data collection

We obtained Ontario antler data from The Foundation for the Recognition of Ontario Wildlife (FROW) for the years 1941 to 2020. FROW is a non-profit organization that manages the records of big game animals in Ontario. Antler measurements for white-tailed deer were voluntarily submitted by hunters to FROW from every county in Ontario, except for Temiskaming. The antler dataset included date of kill (DOK), antler type (typical or atypical), tine numbers and multiple scores following the Boone and Crockett scoring system, which assigns scores based on additive measurements of the antler including tine length, beam circumference, and spread. FROW has a minimum requirement that must be met in order for a white-tailed deer to be entered into the record book (atypical net score 130 and typical net score 120; Foundation for the Recognition of Ontario Wildlife 2015), so we refer to these deer as “record deer” hereafter. Minimum scoring requirements increased in 2006, so all records prior to 2006 below this adjusted score were excluded. Because we were interested in assessing the influence of spatial and temporal variables on records, we removed entries that did not contain a county or DOK. Lack of landcover data dating back pre--1999 resulted in all record deer entries prior to 1997 being removed in the spatial models. We then assessed the influence of a series of environmental and temporal variables on tine number, gross score, net score, symmetry and the number of record deer per county.

Predictor variables in our model included temperature (°C), precipitation (mm), and number of growing degree days (GDD, a measure of accumulated warmth above a threshold needed for plant growth) for each county corresponding to the year a record deer was harvested (Table S1, available in Supporting Information). These variables were obtained from Natural Resources Canada (Hutchinson et al. 2009, Hopkinson et al. 2011, McKenney et al. 2011). We extracted temperature and precipitation data from the months March to August as these are the months when antler growth occurs (Hewitt 2011). For precipitation, we also extracted the months January to March to gauge winter severity through snow fall. Precipitation data were averaged. For temperature, we followed Siegert *et al*. (2013) and calculated the yearly temperature difference by subtracting each county’s average temperature over the time series from the temperature for each individual year. The temperature and precipitation data for one year prior to the DOK and two years prior to the DOK were also extracted.

ArcGIS pro (version 3.1.3;(Johnston and Environmental Systems Research Institute 2004), was used to extract landcover data. We used two land cover layers: the Ontario Land Cover Compilation v.2.0 (Land Information Ontario 2023) which captured the years 1999 to 2011 and Ontario Land Cover Version 1.0 (Ontario Land Cover Version 1.0 - Overview n.d.) which captured the years 2015 to 2020. Both land cover layers are raster datasets with a 15-metre spatial resolution and a minimum mapping unit of 0.5 hectares. Depending on their DOK, the record deer were assigned to a landcover dataset (Table S1). For the years not captured by the landcover databases (2012-2014), whichever layer encompassed the year closest to the DOK was used. Reclassification of the land cover categories within these databases was completed to ensure consistency between land cover types: reclassified categories were; agriculture, infrastructure, deciduous forest, coniferous forest, mixed forest, other forest, rangeland, water, wetlands, and other landcover types (Fig. 1, Table S2). The proportion of each landcover type per county was calculated (Table 1), and “other landcover” was dropped in order to interpret the relationship between the response variable and the proportion covariates (see Valle *et al*., 2024).

**Figure 1.**
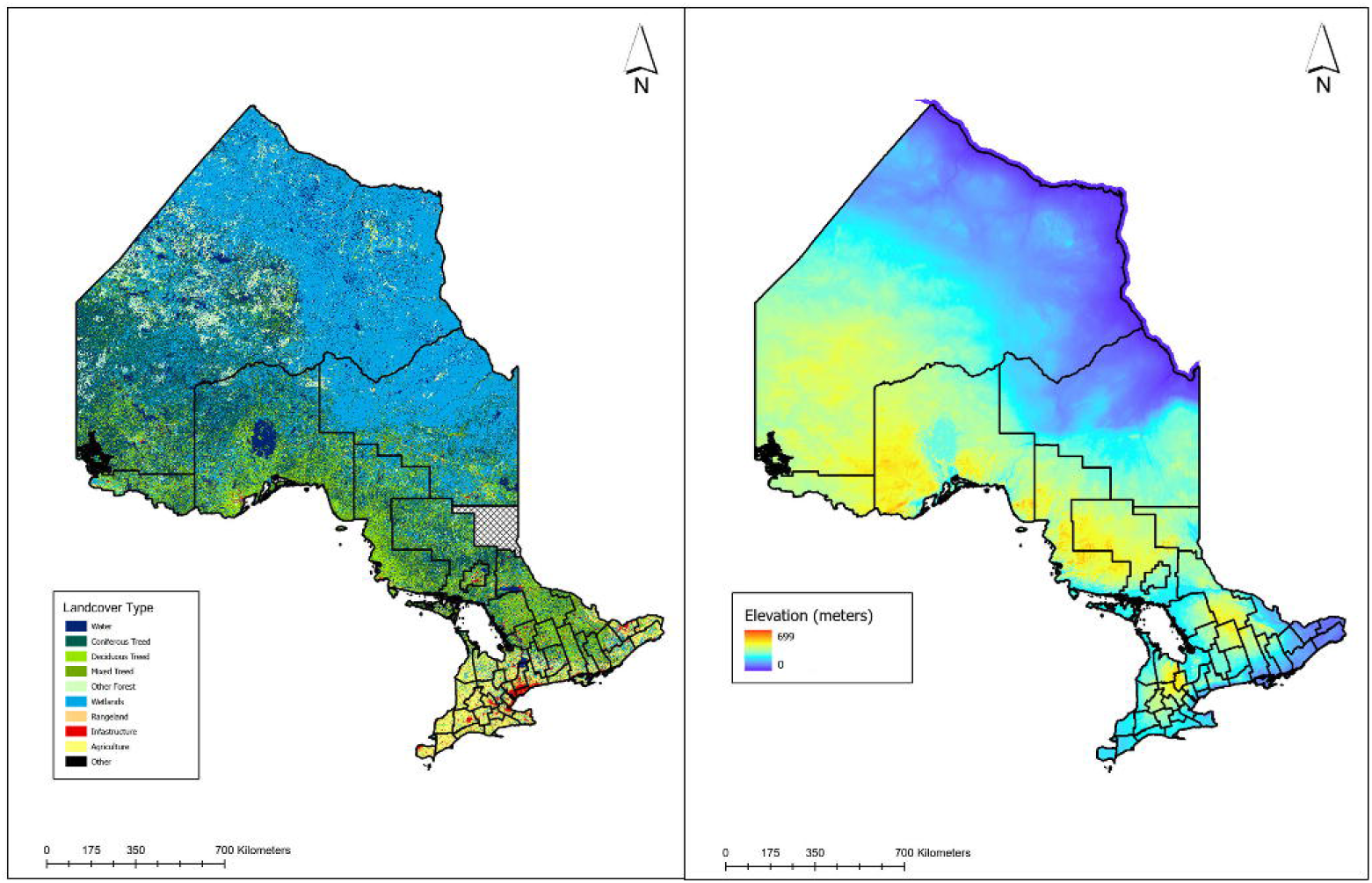
Approximate Ontario land cover distribution (Left) and elevation (Right) divided by county.

**Table 1.** Percentage of land cover types in Ontario, 1999 – 2011 vs 2015 – 2020.

The final three predictor variables used were average county elevation (meters; TessaDEM v1.2 2024), county size (km^2^) and deer abundance (Table S1). Average county elevation and county size are fixed variables, with the values used per county staying constant over the study period. A relative index of white-tailed deer abundance per county was generated using data from provincial hunter surveys (Government of Ontario 2025). Each year, the Ontario Ministry of Natural Resources collects information online or by phone through mandatory hunter surveys on hunting activities. As part of this survey, hunters are asked to report the number of deer they saw during their hunt and the total number of days they hunted, providing a metric of the deer seen per day (Government of Ontario 2025). These values were averaged across all available years (2008-2023) in each county to provide a single estimate per county.

### Statistical analysis

The following predictor variables were standardized (centered to a mean of zero and scaled to a standard deviation of one) prior to analysis: average county elevation, precipitation (winter and antler growing season - March to August), estimated deer abundance, and GDD. Temperature predictors were not standardized because they were already transformed to reflect deviations from long-term county-level averages. We assessed pairwise correlations among the variables, removing those with a correlation greater than 0.70 or less than -0.70 (i.e. Dormann et al. 2013). Landcover variables were excluded from the correlation assessment among themselves since they are proportion data, therefore, changes in one necessarily correspond to changes in others, making high correlations expected and not indicative of problematic collinearity (Valle et al. 2024). However, we did assess correlations between landcover variables and other covariates. We then fit a series of models to each response variable. All spatial models were fitted using the S.CARmultilevel function from the CARBayes package in R ( v.6.1.1Lee, 2013, R Core Team 2023). These models capture spatial autocorrelation by incorporating spatial random effects modeled with a Leroux conditional autoregressive (CAR) prior, where neighboring counties are defined by shared borders. Model fitting occurred within a Bayesian framework using Markov chain Monte Carlo (MCMC) simulation with default priors applied to all model parameters. All model specifications, including response variables and full lists of predictors, can be found in Table S3, available in supporting information.

To assess the number of record deer harvested in each county we used a Poisson regression with a log link and a county area offset to translate the analysis into one of rates of record deer per area. We summed the total number of record deer harvested per county across all years, so DOK was not considered when assigning temperature and precipitation in this model. Instead, average values for the temperature and precipitation of each county over the specified time span were used. The averaging of these environmental variables allowed us to observe relationships between the environment and the spatial distribution of record deer harvests across Ontario counties over the entire study period.

To evaluate antler characteristics, we used DOK when assigning temperature, precipitation and landcover values which allowed us to observe spatial-temporal relationships at the individual level. Each model used one of three distributions and their standard link functions: Poisson regression model with a log link to assess the number of antler tines, a binomial regression model with a logistic link (logit) to assess antler symmetry (symmetrical = 0, asymmetrical = 1), with asymmetry defined as unequal tine counts between antlers rather than the broader Boone and Crockett “non-typical” classification, and gaussian regression models with identity links to assess antler score (Net and Gross). Antler score was also log-transformed to normalize the distribution and ensure the assumptions of the model were met.

For CAR models, we evaluated parameter effects based on the probability of direction (pd) which provided the probability each covariate was either positively or negatively related to antler characteristics (Makowski et al. 2019). Parameters with a probability ≥ 97.5% were considered to have a high probability of affecting the antler trait. Parameters with a probability of 95 – 97.5% were considered to have a moderate probability of affecting the antler trait, and parameters with a probability of 90 – 95% were considered to have a weak, but present, probability of affecting the antler trait. All probabilities below 90% were considered to have no effect (Lohr et al. 2020, Dias et al. 2025). Spatial autocorrelation (ρ) was interpreted such that values near 0 indicated spatial independence and values near 1 indicated strong spatial dependence, so, ρ was considered high if > 0.7, moderate if 0.3–0.7, and low if < 0.3. Spatial variance (τ²) and residual variance (σ²) were considered high if values approached or exceeded 1 and low if around or below 0.01, based on the Inverse-Gamma (1, 0.01) priors, which assume most variation is near zero unless the data strongly indicates otherwise (Lee 2013).

Finally, standard generalized linear models (GLMs) from the base r package (R Core Team 2023) were used to assess temporal trends in antler score and tine number, with year as the predictor variable. Harvest data was sparse pre 1980 and so we used only entries from 1980 onward for these GLMs. No additional offsets or weights were applied to the models. Model assumptions were assessed using standard analyses, including assessing normality and homoscedasticity of residuals for the Gaussian models, and overdispersion for Poisson and Binomial models. It should also be noted that this data was hunter-submitted and therefore likely to be non-random and influenced by variation in hunting effort or harvest efficiency, which was not explicitly captured in our models.

## RESULTS

The final dataset after cleaning consisted of 2,413 entries between 1997 and 2020, and 3,072 entries between 1980 and 2020. Number of growing days was removed due to its high correlation with temperature (Fig S1, available in Supporting Information). Temperature was kept as it directly reflects seasonal climate variation and is easier to interpret. Deer abundance was also removed due to its high correlation with agricultural landcover (Fig S1, available in Supporting Information). Residuals from gaussian models were normal, and no overdispersion was detected in binomial or Poisson models.

Average temperature had a strong probability of having a positive effect on the number of record deer per county (Fig. 2; β = 1.24, pd = 99.9%), with years 1 °C warmer than the county average associated with 3.5 times more record deer, corresponding to a 1.24 increase on the log scale. Elevation showed a weak probability of having a positive effect (Fig. 2;β = 0.31, pd = 94.3%), with a 0.35% increase in the expected number of record deer per 1 m increase in elevation. This model had a strong τ² (0.98) and moderate ρ (0.46). For tine number, the temperature one year prior to the year of harvest had a strong probability of having a positive effect (Fig 2; β = 0.02, pd = 99.1%), with a 1 °C increase in the average temperature during the year prior to harvest associated with 2% more tines. The temperature of the year of harvest and the % of Rangeland per county both had weak probability of having positive effects (Fig 2 & 3; β = 0.01, pd = 91.1% and β = 0.94, pd = 92.2%), suggesting that years 1 °C warmer than the county average and counties with a higher percentage of rangeland were associated with 1% and 156% (2.6-fold) more tines, respectively. This model had a low τ² (0.003) and moderate ρ (0.63). For antler symmetry, precipitation during antler growth two years prior to the year of harvest had a strong probability of having a negative effect (Fig 2; β = -0.12, pd = 98.6%), corresponding to a 0.85% decrease in the odds of asymmetrically antlered deer per 1 mm increase in precipitation, while the temperature of the year of harvest had a moderate probability of having a positive effect (Fig 2; β = 0.10, pd = 95.6%), corresponding to an 11% increase in the odds of asymmetrically antlered deer per 1□°C warmer year. This model had a low τ² (0.03) and moderate ρ (0.38).

**Figure 2.**
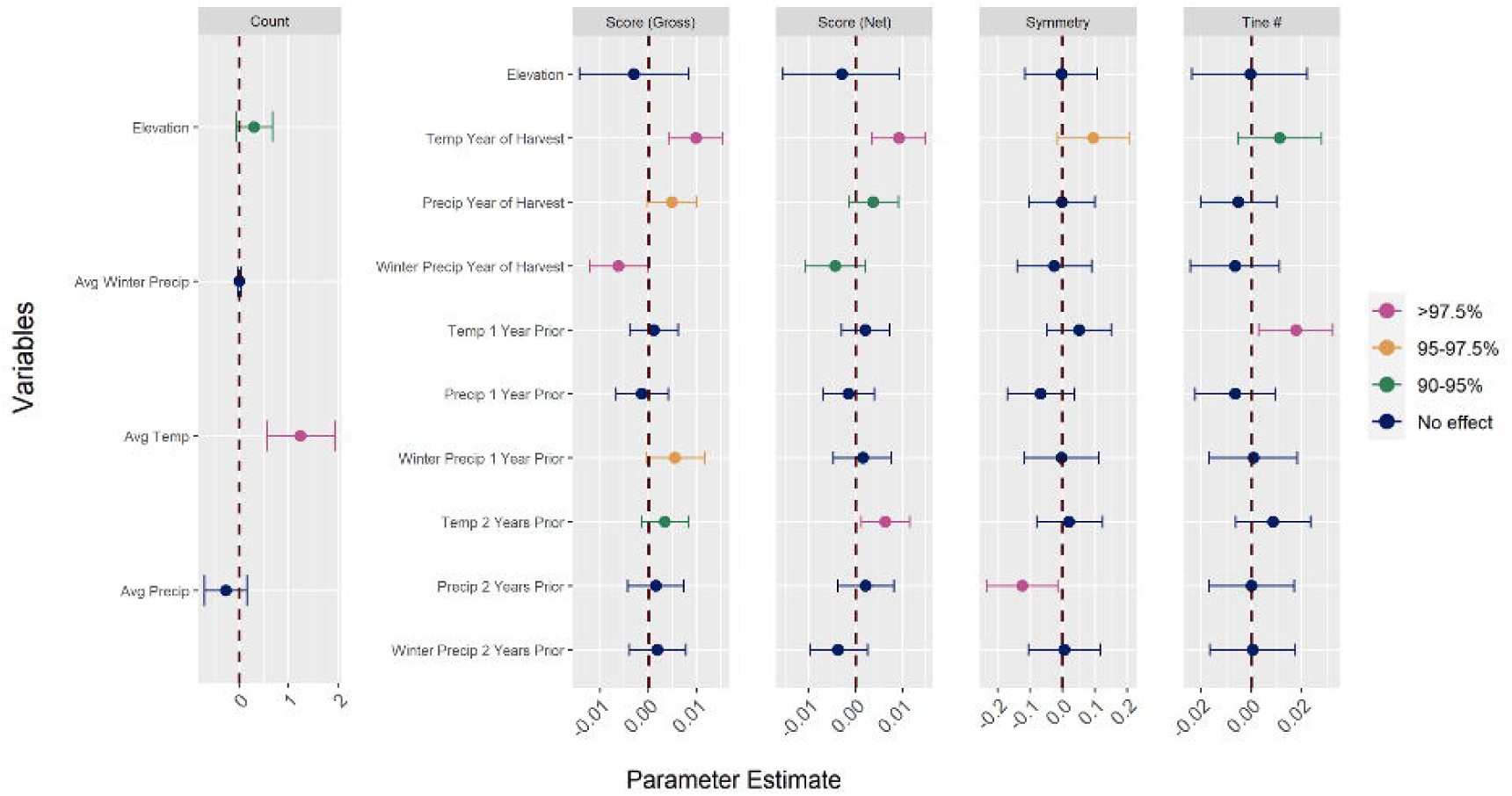
Probability of direction (pd) of elevation, temperature, and precipitation on the response variables. Variables in pink have a high probability affecting the antler trait (97.5%). Variables in Orange have a moderate probability (95 – 97.5%). And Variables in green have a weak, but present probability (90 – 95%).

**Figure 3.**
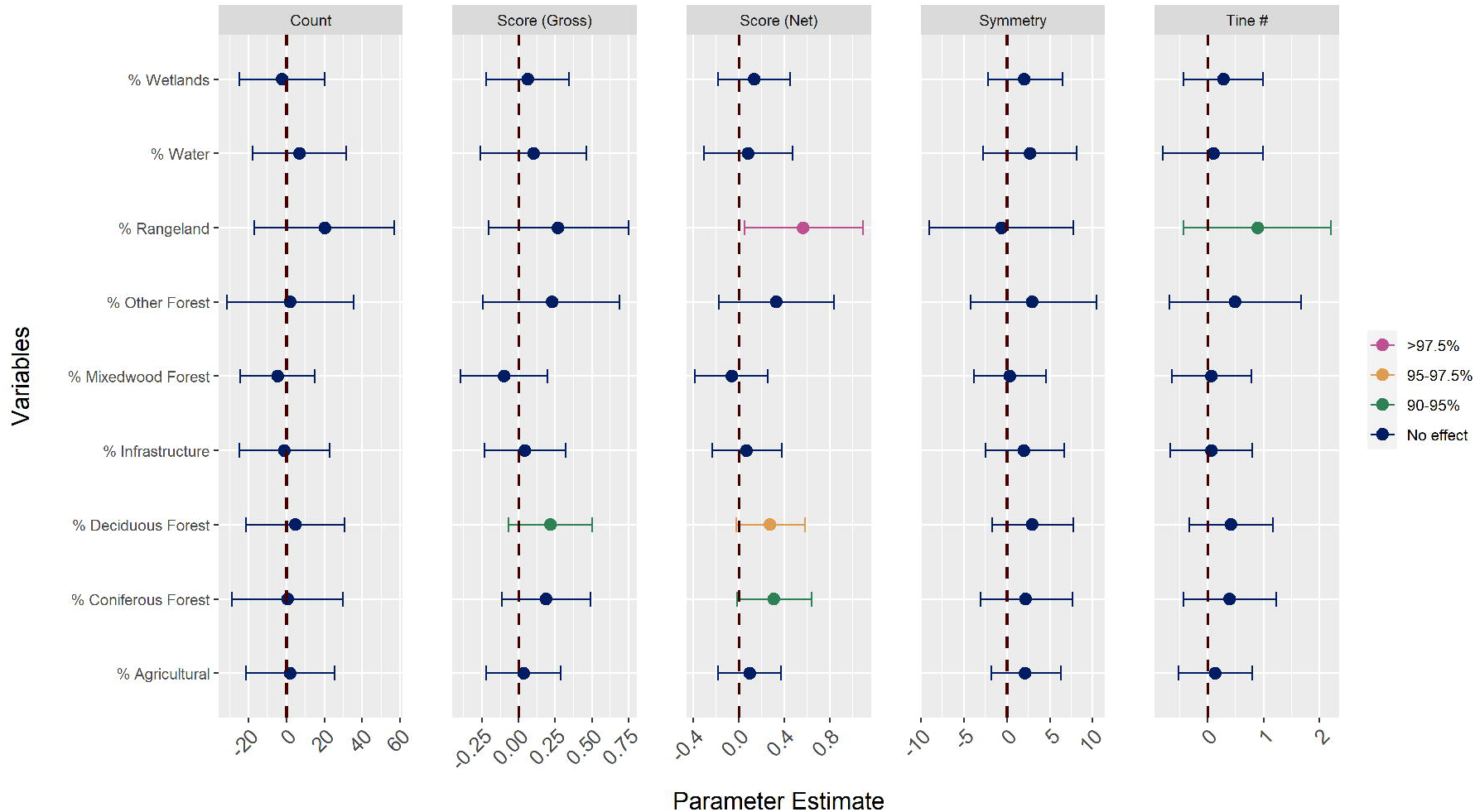
Probability of direction (pd) of landcover proportion on the response variables. Variables in pink have a high probability affecting the antler trait (97.5%). Variables in Orange have a moderate probability (95 – 97.5%). And Variables in green have a weak, but present probability (90 – 95%).

For Gross Score, we found the temperature the year of harvest had a strong probability of having a positive effect (Fig 2; β = 0.01, pd = 100%), with years 1 °C warmer than the county average associated with a 1% increase in Gross Score. Winter precipitation during the year of harvest had a strong probability of having a negative effect (Fig 2; β = -0.01, pd = 97.8%), with a 1 mm increase in winter precipitation associated with a 0.05% decrease in Gross Score. Precipitation during antler growth the year of harvest and precipitation during the winter one year prior to the year of harvest both had moderate probabilities of having positive effects (Fig 2; β = 0.01, pd = 96.8% and β = 0.01, 96.5%, respectively), with a 1 mm increase in precipitation during antler growth the year of harvest being associated with a 0.07% increase in Gross Score and a 0.05% increase in Gross Score for winter precipitation one year prior to harvest. Temperature two years prior to harvest and the % of Deciduous Forest per county then had weak probabilities of positive effects (Fig 2 & 3; β = 0.003, pd = 91.4% and β = 0.22, 93.4%), with 0.3% increase per 1□°C above the county average and 25% increase per 1% more Deciduous Forest, respectively. This model had τ² and ^2^, both of which were relatively low (0.001 and 0.01 respectively), and a moderate ρ (0.70).

For Net Score, temperature the year of harvest and temperature two years prior to the year of harvest had strong probabilities of having a positive effect (Fig 2, β = 0.01, pd = 100%, β = 0.01, pd = 99.2% respectively), with each suggesting a 1% increase in Net Score per 1 °C increase above the county average. The % of rangeland per county also had strong probabilities of having a positive effect (β = 0.58, pd = 98.9%), with a 79% increase in Net Score per 1% more rangeland. The % of Deciduous Forest per county then had a moderate probability of having a positive effect (Fig 3, β = 0.27, pd = 96.0%), with a 31% increase in Net Score per 1% more deciduous forest. The precipitation during antler growth the year of harvest and the % of Coniferous Forest per county both had weak probabilities of positive effects (Fig 2 & 3; β = 0.004, pd = 92.8% and β = 0.28, pd = 94.8%, respectively), with a 1 mm increase in precipitation during antler growth the year of harvest being associated with a 0.03% increase in Net Score and a 1% increase in coniferous forest being associated with a 32% increase in Net Score. The precipitation during the winter the year of harvest then had a probability of having a weak but negative effect (Fig 2; β = -0.004, pd = 91.1%), with a 1 mm increase in precipitation during the winter the year of harvest being associated with 0.02% decrease in Net Score. τ² and ^2^ were relatively low (0.002 and 0.01 respectively), and ρ was moderately high (β =0.74). Finally, there was no temporal trend in size and tine number over time (Table S4, available in Supporting Information).

## DISCUSSION

We investigated spatial drivers and temporal patterns of harvested record deer in Ontario, Canada. We observed that increases in temperature during the months that antler growth is occurring (March-August) positively influenced all antler characteristics (Fig 2), and that rangeland and forest are key landscape components, the amounts of which are correlated to overall greater antler size. These patterns are consistent with our hypothesis that conditions providing abundant forage and adequate cover will promote antler growth. Higher winter precipitation had negative effects, consistent with our prediction that conditions increasing energetic demands, such as deep snow and limited forage, would hinder antler growth. Interestingly, no change in antler size was detected over time in Ontario (Fig. 4), which does not support the idea of regular harvest of large-antlered individuals leading to a temporal decline through reductions in age structure. This finding is likely due to the province’s position at the northern edge of the white-tailed deer’s range and a relatively stable population age structure. Historically, harsher winters and limited forage could have constrained antler growth, but ongoing warming due to climate change is likely improving habitat suitability by reducing winter severity and extending growing seasons, which could be helping maintain antler size in Ontario (Lesage et al. 2001, Dawe and Boutin 2016).

**Figure 4.**
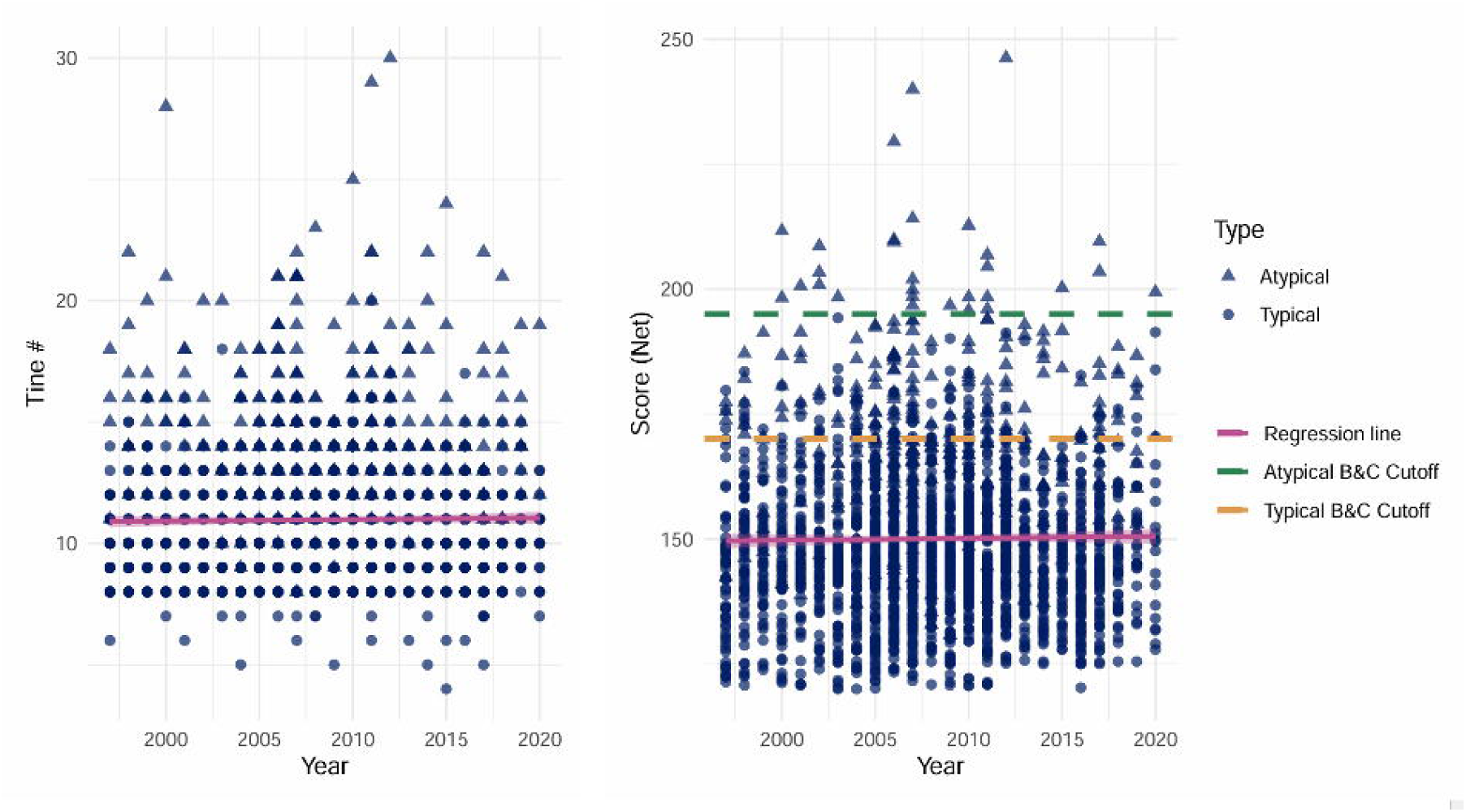
Change in antler score and tine number over time. Cutoffs are representative of Boone and Crockett data, depicting the higher cutoff size for record antlers in other parts of North America.

This data set is representative of adult antlered white-tailed deer within Ontario and although not a random sample, the data captures a large range of variation in antler size across the population (Fig S2, available in Supporting Information) and temporal, spatial and environmental influences should be reflected in this large sample of male white-tailed deer (see also Strickland and Demarais, 2008; Monteith *et al*., 2013; Cain *et al*., 2019). However, because participation in record programs is voluntary and biased toward larger antlered deer, those with smaller antlers are likely underrepresented (LaSharr et al. 2019). As a result, the dataset might not fully represent population wide variation in antler size, but relative patterns and relationships with environmental variables that are still informative and can help to understand the factors influencing antler development across Ontario (LaSharr et al. 2019). It should also be noted that, although some effect sizes presented are small, they nonetheless represent measurable changes that will be considered in the interpretation of this study.

To account for spatial dependence in our data and better understand regional patterns, we used CAR models, which revealed moderate to high spatial autocorrelation (ρ) in all models, revealing that nearby counties tend to have similar conditions (e.g., temperature). However, these values also indicate that some unaccounted-for spatial covariates are still present but not explicitly modeled. Possible missing covariates worth further exploration include land productivity, which affects forage availability, landcover structure that shapes shelter and feeding opportunities, and the distribution of minerals like phosphorus and calcium, which is influenced by vegetation type and tissue age and affects bone and antler development (Grasman and Hellgren 1993, Cain et al. 2019, Turner et al. 2025), though the appropriate GIS layers currently do not exist for Ontario. We also observed low τ² in models assessing antler characteristics indicating that, although the predictors often have similar conditions in neighboring counties, antler size does not exhibit strong spatial structure. However, a strong τ² when assessing the count of record deer per county does suggest that counties with higher numbers of record deer are more likely to have neighboring counties with similarly high counts, likely reflective of similar forage resources and variation in regional hunting interest and behaviour. Notably, hunter behaviour and success which will vary with the amount of roads, dwellings, and with accessibility, while regional differences in the number of hunters and frequency of hunting might also contribute to these spatial patterns (Kilgo et al. 1998, Bonnot et al. 2013).

### Environmental drivers of record antlers

Temperature was the most consistent environmental driver of antler traits: higher temperature was related to an increase in antler size metrics in all models (Fig 2). Warmer temperatures have been linked to larger antlers and greater reproductive success in white-tailed deer and other Cervidae (Clements et al. 2010, Simard et al. 2010). Increased plant growth from warmer conditions means more forage availability for the deer, which will supply the nutrients and energy needed for antler growth (French et al. 1956, Hatfield and Prueger 2015). Here, the unique annual regrowth of antlers makes them strongly affected by yearly environmental changes (Foley et al. 2012). When looking at tine number and score, the temperature one year prior to harvest and two years prior to harvest respectively, also had positive effects on antler size and shape (Fig. 2). Antler growth depends on the accumulation of resources stored in the animal’s body (i.e. mineral and fat reserves) (French et al. 1956, Lesage et al. 2001). Warmer conditions that promote forage growth in prior years can enhance fat reserves, which are then used in subsequent antler cycles when these nutrients is needed (Hewitt 2011).

Similar inference can be made from the influence of precipitation during antler growth the year of harvest on antler size. Wetter conditions can lead to increased plant growth which could again affect the annual regrowth of antlers (Foley et al. 2012, Scharwies and Dinneny 2019). Conversely, we saw negative effects from precipitation during the winter the year of harvest on antler size (Fig 2). More snow in winter can make it more difficult for the deer to move around and find forage, which causes them to congregate in deer yards (Government of Ontario 2019*b*). Deer yards are an area made up of coniferous trees that have a large canopy for blocking excessive snowfall (Government of Ontario 2019*b*). With deer being confined to these yards deer density and the availability of forage for each individual animal decreases (Patterson and Power 2002). This could then put individuals with poor body conditions at a disadvantage and minimal fat reserves when antler regrowth commences (Lesage et al. 2001). Future work could strengthen this finding by incorporating direct measurements of snow depth, snow cover duration, or broader indices such as the Winter Severity Index (WSI) to capture more details on the energetic challenges white-tailed deer face during the winter months.

Rangeland and forest landcover had positive effects on antler characteristics, meaning that as the percentages of these types of landcover in each county rose so too did antler score and tine number (Fig 5). Areas combining forest with regions that provide abundant forage have been associated with higher harvests of record deer, likely due to greater amounts of high-contrast edges, which create habitat interfaces that offer the resources necessary for male deer to produce large antlers (Cain et al. 2019). In our case, forests provide cover, which block excessive snowfall, and contain the northern deer’s primary winter food source, browse (Hewitt 2011), and rangeland provides a more variable and nutritious food sources such as forbs, while also allowing for greater visibility of large antlered bucks from further away, potentially increasing hunter success or improving the effectiveness of dominance displays and mate attraction (Gilbert et al. 2006, Hewitt 2011, Houde et al. 2020). Collectively, these habitat interfaces are preferred by white-tailed deer as it allows them to have a larger variety of food sources throughout the year and provides both more space for movement among resources (rangeland) while still having adequate cover from predators and severe winters (forest) (Voigt et al. 1997, Bartel et al. 2025).

**Figure 5.**
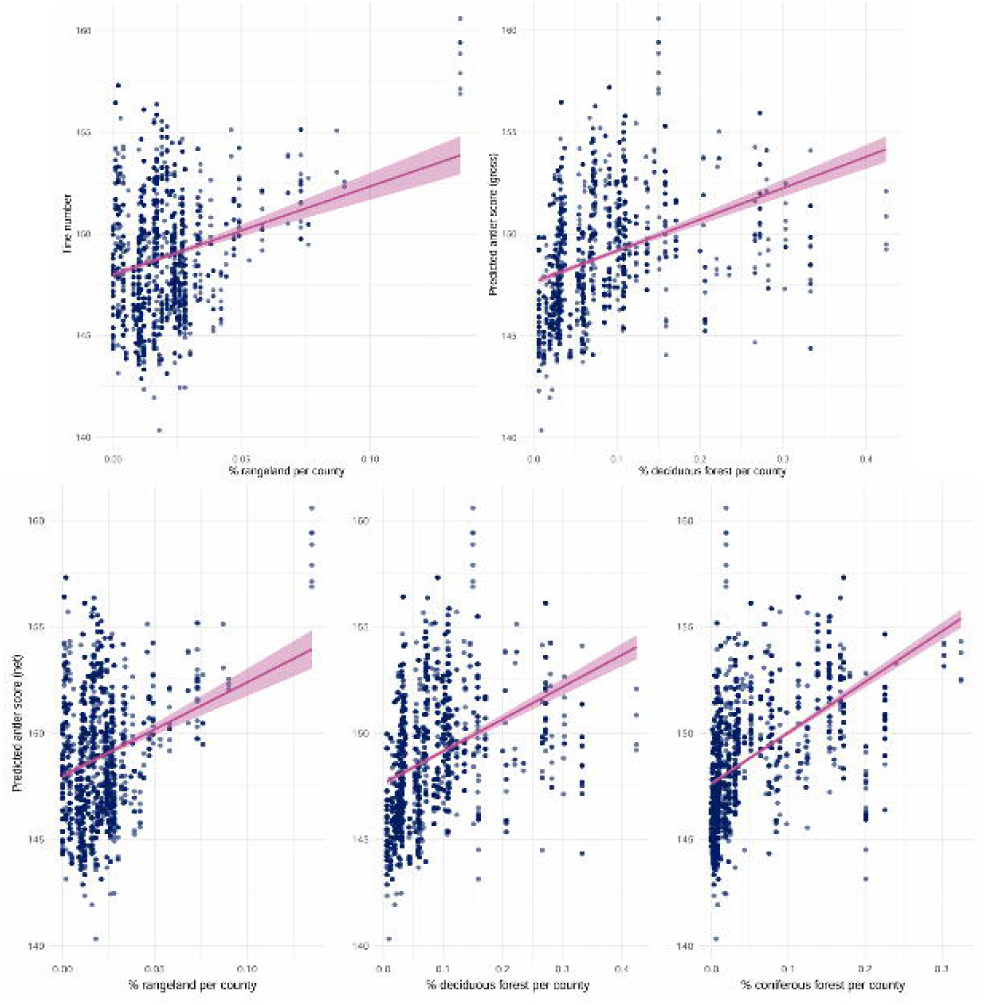
Percent of significant landcover per predicted antler score and tine number.

### Number of harvested deer and antler symmetry

We observed that increased average temperatures were associated with more record white-tailed deer being harvested in a given Ontario county with years 1 °C warmer than the county average associated with 3.5 times more record-scoring deer in a given county. Similar to our assessment of this trend seen in our antler size and configuration models, warmer temperatures result in more forage at the northern edge of the white-tailed deer’s range and thus more energy is able to be allocated to individual deer in a county, presumably allowing more individuals to reach record deer status. The number of record deer harvested was not significantly affected by any landcover covariates (Fig 3); so, while landcover type positively impacts antler size, the overall effect is minor such that it does not increase the net number of large animals harvested in a given year. Curiously, record deer numbers were positively affected by elevation (Fig. 2). It is tempting to suggest this being reflective of mature deer finding refuge from human activity via fewer roadways at these elevations. Less trafficked zones allow deer to move more freely and access critical resources with minimal human interference (Hamilton et al. 2024) and behaviourally deer learn to avoid humans (Kilgo et al. 1998, Marantz et al. 2016). However, the elevational range is small in the province (0 m – 700 m) and obfuscated when averaged across the county as done here (66 m – 415 m), and thus finer scale analysis is warranted to better understand this effect.

The symmetry of antlers appears to be impacted by precipitation, with higher precipitation levels two years prior to antler growth resulting in more symmetrically antlered deer. Since white-tailed deer can develop antler asymmetry when exposed to stressors such as environmental conditions, disease, or injury that affect energy allocation before and during antler growth (Marburger et al. 1972, Goss 1983, Wild et al. 2022), the mechanism by which precipitation two years prior impacts symmetry is still unclear. However, there is the potential for lagged environmental effects on antler morphology which has been observed in other studies, with Strickland et al. (2020) demonstrating that conditions experienced by male white-tailed deer two years prior were associated with variation in antler mass. Likewise, Patterson and Power (2002) suggested that antler beam size was affected by lagged climatic effects through influences on habit and population density. Symmetry has also been seen to decrease as trait size increases (De Coster et al. 2013), which we observed in our data set (Fig S3, available in Supporting Information); however, we see no strong relationship between precipitation and antler size. It is thus still conceivable that environmental conditions might have long-term impacts on symmetry, even for antlers that are shed annually.

### Temporal patterns of record deer

Over the last 40 years, there has been no significant change in antler size (or score) and the tine number of harvested record white-tailed deer in Ontario (Table S4). This contrasts the findings of Monteith et al. (2013) who reported a negative trend across North America in antler size in white-tailed deer from 1950 to 2008, based on trophy records of mature males submitted to the Boone and Crockett Club’s *Records of North American Big Game* (Boone and Crockett Club 2025). These negative trends were hypothesized to be a combination of harvest-induced reduction in age structure, sociological effects and density dependence (Monteith et al. 2013). Our finding suggests that antler trends in white-tailed deer are region specific and vary depending on the environment and pressures present.

In Ontario, the number of active hunters has stayed relatively consistent since 2008 while the number of records submitted to FROW has declined (Fig 6). This could be indicative of a possible shift in hunting priorities (Monteith et al. 2013), but multiple severe winters in 2007/08 and 2010/11, have coincided with apparent population declines (Ontario Outdoors Magazine Game Forecast 2009, 2011) which would explain a decrease in entries. Still, the number of white-tailed deer in Ontario is relatively low especially compared to other regions of North America where population numbers are substantially higher. For example, Michigan with similar environmental conditions to Ontario and an area of ∼ 250,493 km^2^ has a population of ∼2 million white-tailed deer, compared to Ontario with an area of over 1 million km^2^ and a white-tailed deer population of ∼400 thousand (Mason 2016, Conner 2021). This lower density is likely due to Ontario being the northern edge of the white-tailed deer’s range, where limited suitable habitat likely has more influence on antlers than harvest pressure or hunting regulations. While Ontario’s hunting regulations could result in lower overall harvest rates compared to other regions, it is not clear if these restrictions make any substantial difference in population dynamics or antler development. Instead, environmental factors associated with climate and habitat likely have a greater influence on population density and antler growth in the province.

**Figure 6.**
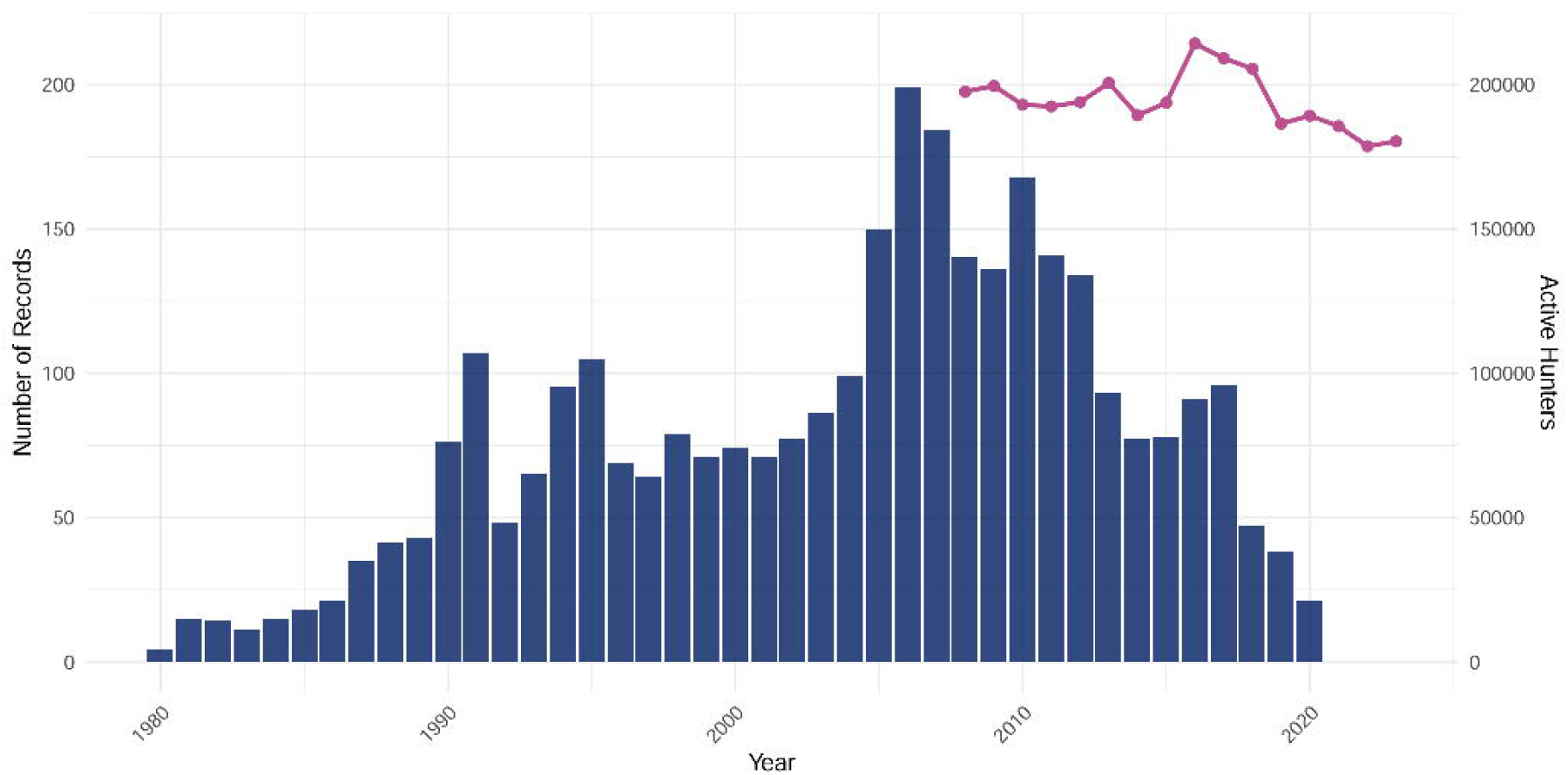
Number of records submitted to the Foundation for the Recognition of Ontario Wildlife (FROW) from 1997 to 2020 (blue) and Number of active hunters from 2008 to 2023 (pink).

In some parts of North America, warmer climates lead to longer growing seasons allowing white-tailed deer to have more time to access forage and accumulate resources, supporting antler development allowing them to reach their maximum physical size during antler growth (Mysterud et al. 2005, Sontheimer et al. 2024). In contrast, Ontario’s relatively shorter growing seasons and colder winters limit access to nutritious food and deplete body reserves, which could reduce the amount resources deer can allocate to antler growth in the subsequent spring; this limitation could mean there is still room for growth. Our models indicate that warmer temperatures in Ontario are positively influencing antler development, likely because milder winters and longer growing seasons improve food availability. Combined these conditions could be helping bolster antler size despite pressures that are causing declines elsewhere (e.g. Monteith et al. 2013).

## Supporting information

All other Supplemental Figures and Tables

## MANAGEMENT IMPLICATIONS

Our findings highlight how variation in habitat conditions across Ontario counties are related to differences in antler size and tine configuration as well as record white-tailed deer distributions. This emphasizes the importance of interpreting phenotypic trends relative to specific areas rather than generalizing across entire populations. Because environmental factors such as land cover composition and forage availability influence antler growth, where larger antlered bucks are desired, management practices could focus on maintaining or enhancing the quality of white-tailed deer habitats in Ontario, particularly in areas where forage is limited. Efforts such as sustaining forested habitat interfaces and managing deer densities to balance browse pressure could help improve nutritional conditions that support antler development.

With no significant temporal trend in antler size, the current harvest practices in Ontario are likely effective as it relates to maintaining antler size, but continued monitoring could be carried out to ensure that population dynamics remain stable. The long-term collection of harvest and antler data will be critical for tracking population health and evaluating the effects of management practices. Finally, by communicating these findings, hunters can better manage their expectations and recognize aspects of climate that could cause certain regions and years to have higher large-antlered white-tailed deer concentrations.

## ACKNOWLEDGMENTS

This work was supported by NSERC Discovery Grant (ABAS; 410 SDC); Canadian Foundation for Innovation (ABAS): John R. Evans Leaders Fund (ABAS) and the Ontario Early Researcher Award (ABAS). Trent University is located on the traditional territory of the Michi Saagiig Anishnaabeg. We thank 4 anonymous reviewers for their helpful comments.

## ETHICS STATEMENT

No approval of research ethics committees was required to accomplish the goals of this study.

## DATA AVAILABILITY

The antler data that support the findings of this study are available from The Foundation for the Recognition of Ontario Wildlife, Ontario, Canada and were acquired under research agreements for this study. All other model covariates are provided.

**AAPPENDIX A. SUPPLEMENTAL**

