## Supplementary material for "Environmental and temporal factors affecting record white-tailed deer antler characteristics in Ontario, Canada": All other Supplemental Figures and Tables

January 6^th^, 2026

**Table S1:** Raw climate and land cover variables

**Table S2:** Landcover classification organization

| **Coarse**  **Classification** | **1999** | **2015** |
| --- | --- | --- |
| **Water** | Clear Open Water, Turbid Water | Water |
| **Other** | Shoreline, Disturbance, Cliff and Talus,  Alvar, Sand Barren and Dune, Sand, Gravel,  Mine Tailings, Extraction, Bedrock | Alvar, Dune, Barren |
| **Wetlands** | Mudflats, Marsh, Swamp, Fen, Bog | Coniferous treed swamp,  Mixed wood treed swamp,  Deciduous treed swamp,  Transitional treed swamp,  Thicket swamp, Bog, Fen, Marsh |
| **Other Forest** | Heath, Sparse Treed, Treed Upland,  Plantations - Treed Cultivated, Hedge Rows,  Tallgrass Woodland | Sparse treed, Transitional Forest,  Hedge row |
| **Deciduous Forest** | Deciduous Treed | Deciduous forest |
| **Mixed Forest** | Mixed Treed | Mixed wood forest |
| **Coniferous Forest** | Coniferous Treed | Coniferous forest |
| **Rangeland** | Open Tallgrass Prairie,  Tallgrass Savannah | Prairie, Savanah, Meadow, Shrubland |
| **Infrastructure** | Community, Infrastructure | Built up area-pervious, Anthropogenic,  Transportation |
| **Agriculture** | Agricultural and  Undifferentiated Rural Land Use | Cropland, Hay-pasture |

**Table S3:** Predictor variables used in each model

| ***Response*** | ***Predictors*** |
| --- | --- |
| *Record deer*  *count* | *Avg. elev. + avg. temp. + avg. prec. + avg. prec. winter + avg. # growing days + % water*  *+ % dec. forest + % conif. forest + % mixed wood forest + % other forest + % rangeland*  *+ % wetlands + % infrastructure + % agriculture + abundance + offset(km²)* |
| *Antler tine #* | *Avg. elev. + Δ temp + avg. prec. + avg. prec. winter + # growing days + Δ temp (1 yr prior)*  *+ avg. prec. (1 yr prior) + avg. prec. winter (1 yr prior) + # growing days (1 yr prior)*  *+ Δ temp (2 yrs prior) + avg. prec. (2 yrs prior) + avg. prec. winter (2 yrs prior)*  *+ # growing days (2 yrs prior) + % water + % dec. forest + % conif. forest + % mixed wood forest*  *+ % other forest + % rangeland + % wetlands + % infrastructure + % agriculture + abundance* |
| *Antler*  *symmetry (0/1)* | *Avg. elev. + Δ temp + avg. prec. + avg. prec. winter + # growing days + Δ temp (1 yr prior)*  *+ avg. prec. (1 yr prior) + avg. prec. winter (1 yr prior) + # growing days (1 yr prior)*  *+ Δ temp (2 yrs prior) + avg. prec. (2 yrs prior) + avg. prec. winter (2 yrs prior)*  *+ # growing days (2 yrs prior) + % water + % dec. forest + % conif. forest + % mixed wood forest*  *+ % other forest + % rangeland + % wetlands + % infrastructure + % agriculture + abundance* |
| *Score*  *(gross/net)* | *Avg. elev. + Δ temp + avg. prec. + avg. prec. winter + # growing days + Δ temp (1 yr prior)*  *+ avg. prec. (1 yr prior) + avg. prec. winter (1 yr prior) + # growing days (1 yr prior)*  *+ Δ temp (2 yrs prior) + avg. prec. (2 yrs prior) + avg. prec. winter (2 yrs prior)*  *+ # growing days (2 yrs prior) + % water + % dec. forest + % conif. forest + % mixed wood forest*  *+ % other forest + % rangeland + % wetlands + % infrastructure + % agriculture + abundance* |
| *Score and*  *Antler Tine #*  *(Temporal)* | *Year* |

**Table S4:** Change in Ontario’s white-tailed deer antler characteristics over time (1980-2022).

| **Model** | **Coefficient** | **p-value** | **β** |
| --- | --- | --- | --- |
| Tine number | Year | 0.594 | 0.001 |
| Gross score | Year | 0.501 | 0.039 |
| Net score | Year | 0.997 | -2.462 e-04 |

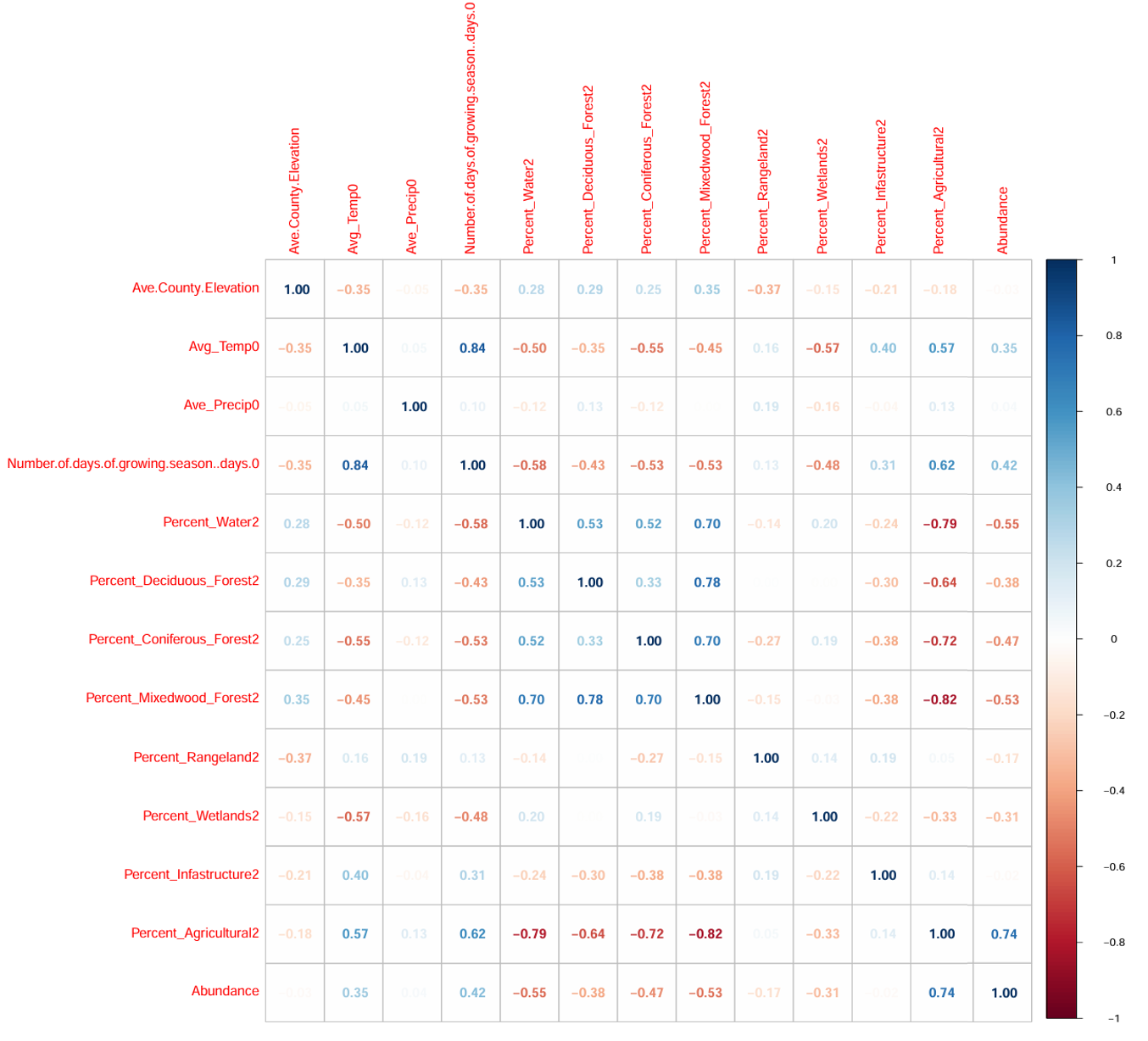

**Fig S1:** Correlation matrix showing relationships among predictor variables used in spatial models.

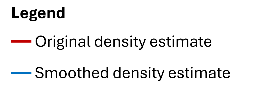

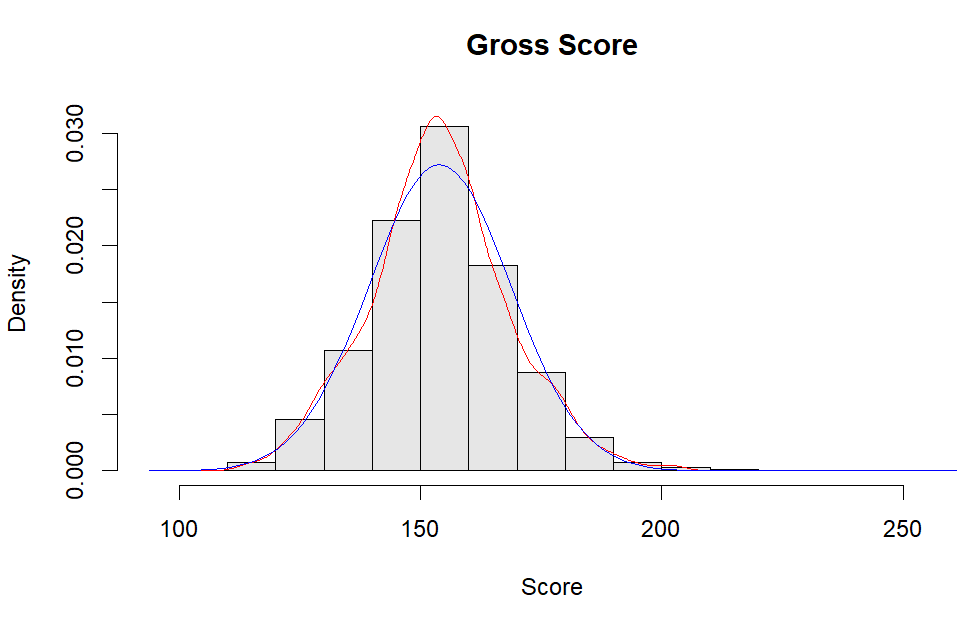

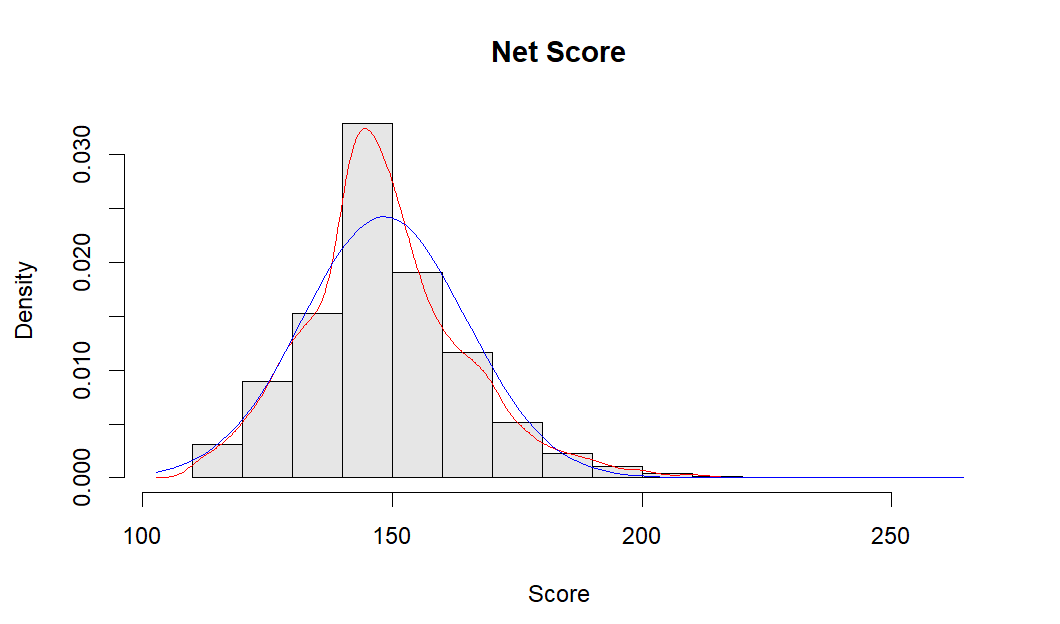

**Fig S2:** Bell curve score distribution. Red line: the estimate using the default bandwidth, capturing more detail in the distribution. Blue line: the estimate using a larger bandwidth to produce a smoother curve

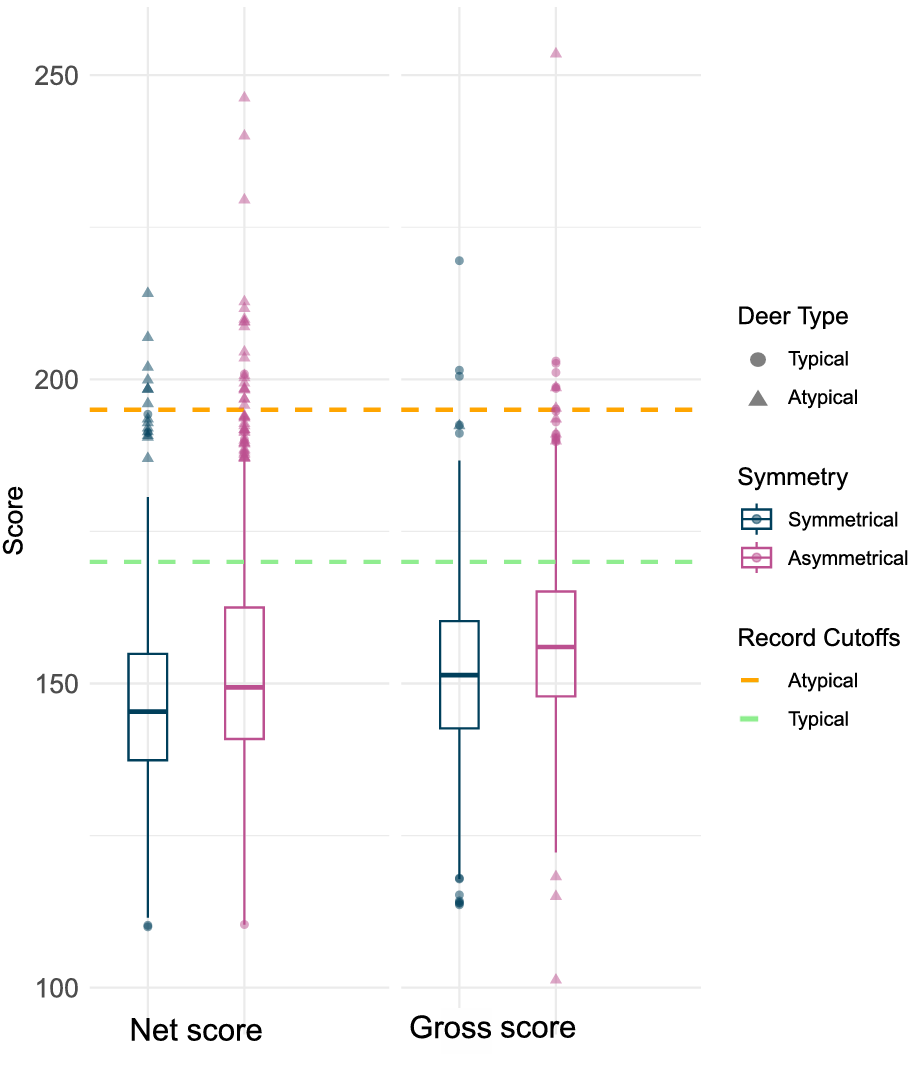

**Fig S3.** Score of symmetrical vs asymmetrical antlered record deer
